# Anti-psychotic drugs act synergistically in combination with antifungal drugs to inhibit drug-resistant *Cryptococcus* and *Candida* species

**DOI:** 10.64898/2025.12.31.697249

**Authors:** Magali Ayala, Alifia Jakamartana, Rodrigo Mamede Dos Santos Costa, Joseph M. Bednarek, Christian T. Moreau, Cheuk-Kiu Eric Cheung, Jessica C. S. Brown

**Affiliations:** School of Biological Sciences, University of Utah, Salt Lake City, UT 84112

## Abstract

Systemic antifungal infections cause an estimated 3.8 million deaths annually, approximately 10% of which are caused by drug-resistant infections. With only five classes of antifungal drugs, treatment options are limited. Here we explore synergistic drug combinations – when the efficacy of two drugs combined is greater than expected based on the sum of each individual drug’s efficacy – to improve treatment of drug-resistant *Cryptococcus neoformans* and *Candida* species. Chlorpromazine acts synergistically with both amphotericin B and fluconazole against multiple fungal species, including azole-resistant *C. neoformans* and *C. auris*. We then performed a genome-wide knockout mutant screen and found that ESCRT pathway mutants are resistant to chlorpromazine and knockout mutants of genes involved in fatty acid biosynthesis are sensitive. Based on these data, we investigated sterols and fatty acids in chlorpromazine-treated cells and found only minor increases in sterol precursors but a substantial increase in lipid droplet size and decrease lipid droplet number. We conclude that chlorpromazine acts by potentially sequesters lipids, preventing lipolysis and lipid mobilization in response to stress. Together, these data suggest that chlorpromazine and its analogs are potentially promising treatments for systemic fungal infections that act via lipid homeostasis and stress response.

## Introduction

Human interactions with environmental fungi are common but benign: every time we breathe, we inhale fungal spores (1). However, a small subset of species cause severe, systemic disease. These infections have a high mortality rate, causing an estimated 3.8 million deaths annually (2). There are only five available drug classes to treat systemic fungal infections (3): 1) polyenes (4; 5) 2) azoles (6), and which target the plasma membrane sterol ergosterol and its synthesis. 3) Nucleoside analogs target DNA and RNA synthesis (7); 4) echinocandins, which target β-1,3-D-glucan synthase in the cell wall (8); and 5) ibrexafungerp (9) the first member of the triterpenoid class, which acts on the same target as echinocandins. Due to the severity of these diseases, in 2020 the World Health Organization published their first fungal disease priority list (10). *Cryptococcus neoformans*, the primary cause of fungal meningitis and the second most common cause of AIDS-related mortality (11), was one of four critical priority status fungal pathogens. Approximately 152,000 cryptococcal meningitis cases were reported in 2020 with an estimated 112,000 deaths (2). Although *C. neoformans* infections disproportionately affect the immunocompromised, particularly those with low CD4^+^ T cell counts (12; 13). However, non-HIV cryptococcosis cases are growing: new estimates reported 22,280 cases in individuals with no apparent underlying disease (2).

The mortality rate from cryptococcal infection varies from 24-75% with treatment availability (2; 14). In low-income settings, treatment is particularly challenging due to insufficient resources and difficulty managing the disease. The standard therapy for cryptococcal meningitis consists of induction treatment with a single dose of liposomal amphotericin B (polyene class) and two weeks of flucytosine (nucleoside analog) or fluconazole (azole class). This is followed by consolidation (8 weeks) and maintenance (12 months) treatment with fluconazole. When amphotericin B (AmB) is unavailable, the alternative treatment is fluconazole monotherapy or a combination of fluconazole plus flucytosine; both alternatives increase the mortality rate (14). This long treatment duration requires high degree of patient compliance and carries the potential for side effects such as nephrotoxicity, hepatoxicity, and bone marrow changes (7; 15).

Combination therapy using a repurposed drug with an existing antifungal drug is a promising alternative approach to treating systemic fungal infections (16). New drug development is costly and complex (17), whereas repurposing allows off-label usage of drugs approved for other indications, allowing bypass of costly clinical trials. In this study, we are particularly interested in synergistic combinations, when the efficacy of two drugs combined is greater than their individual effects (18). Synergistic drug combinations can increase therapeutic success when monotherapy fails to achieve the desired results. For example, the use of single antifungals such as fluconazole for the treatment of fungal infections is associated with the emergence of antifungal resistance (19). Synergistic drug interactions can prevent development of or combat resistance by targeting two key proteins in either the same pathway or by targeting two separate pathways (20; 21; 22; 23).

Drugs not specifically developed as antifungal drugs have been shown to enhance antifungal potency when used in combination with an antifungal drug. A classic example is calcineurin inhibitors, which act synergistically with azoles and echinocandins (24; 25; 26). Multiple groups previously identified commonly prescribed drugs from a wide range of bioactive categories (anti-infective, antidepressants, antineoplastic, among others) that are synergistic with fluconazole against *C. neoformans* by unknown mechanisms (23; 27; 28; 29; 30). We additionally identified that the antipsychotic drug chlorpromazine, which acts synergistically when combined with fluconazole (27) and AmB against *C. neoformans* and *Candida albicans.* Chlorpromazine is an attractive candidate for fungal meningoencephalitis treatment because it crosses the blood-brain barrier (31) and is widely available and relatively inexpensive. The goal of this study is to identify the molecular mechanism underlying these synergistic interactions.

Chlorpromazine is a commonly prescribed first-generation phenothiazine antipsychotic (31) with amphiphilic properties (32). It is primarily used to treat schizophrenia and bipolar disorder (31), so we hypothesized that it could treat cryptococcal meningitis and other diseases caused by disseminating fungal pathogens. Previous studies showed that chlorpromazine inhibits the growth of pathogenic fungi, but the exact mechanism is not fully understood (33). Furthermore, due to chlorpromazine’s dual hydrophobic and hydrophilic regions, it is thought to interact with phospholipids in membrane bilayers (34). Changes in membrane composition affects membrane fluidity (35), transport processes (36), lipid-protein interactions (37; 38), and cell signaling (39). All these processes influence lipid synthesis (40).

Lipid biosynthesis and function are also common antifungal drug targets. Ergosterol, the fungal analog of mammalian cholesterol is essential in fungi (41). AmB changes membrane fluidity by binding and extracting ergosterol from the plasma membrane in *C. neoformans* (42). AmB also increases membrane permeability and reactive oxygen species (42; 43). Azole class drugs target 14α-demethylase (Erg11) in the ergosterol biosynthesis pathway (44).

In this study, we elucidate the molecular mechanisms of AmB and chlorpromazine combination and their synergistic interaction against the fungal pathogen *C. neoformans*. This combination has been effective against >90% of clinical isolates and demonstrated synergy against *Candida* species (spp.), including clinical isolates. We identify chlorpromazine’s interference with lipid metabolism and its ability to disrupt both lipid droplet formation and its breakdown processes. Together these data demonstrate that disrupting ergosterol homeostasis (with AmB or fluconazole) and inducing large lipid droplet formation (chlorpromazine) act synergistically to inhibit fungal cells.

## Results

### Chlorpromazine acts synergistically with multiple antifungals against *Candida* spp. and *C. neoformans* clinical isolates

We first investigated the breadth of the synergistic interaction between chlorpromazine and the antifungals AmB or fluconazole. We chose multiple clinical *C. neoformans* and *Candida* spp. with varying levels of fluconazole resistance and susceptibility (Table 1) and tested for synergy using checkerboard assays. Synergy is defined as at least a 4-fold reduction in the minimum inhibitory concentration (MIC) of each drug and was calculated using the fractional inhibitory concentration index (FICI), where an FICI of ≤ 0.5 indicates synergy (45). The AmB and chlorpromazine combination acted synergistically against 47 of the 52 *C. neoformans* clinical isolates tested, including 22 of the 24 fluconazole-resistant isolates (**Fig. 1A and Table 2**).

**Figure 1.**
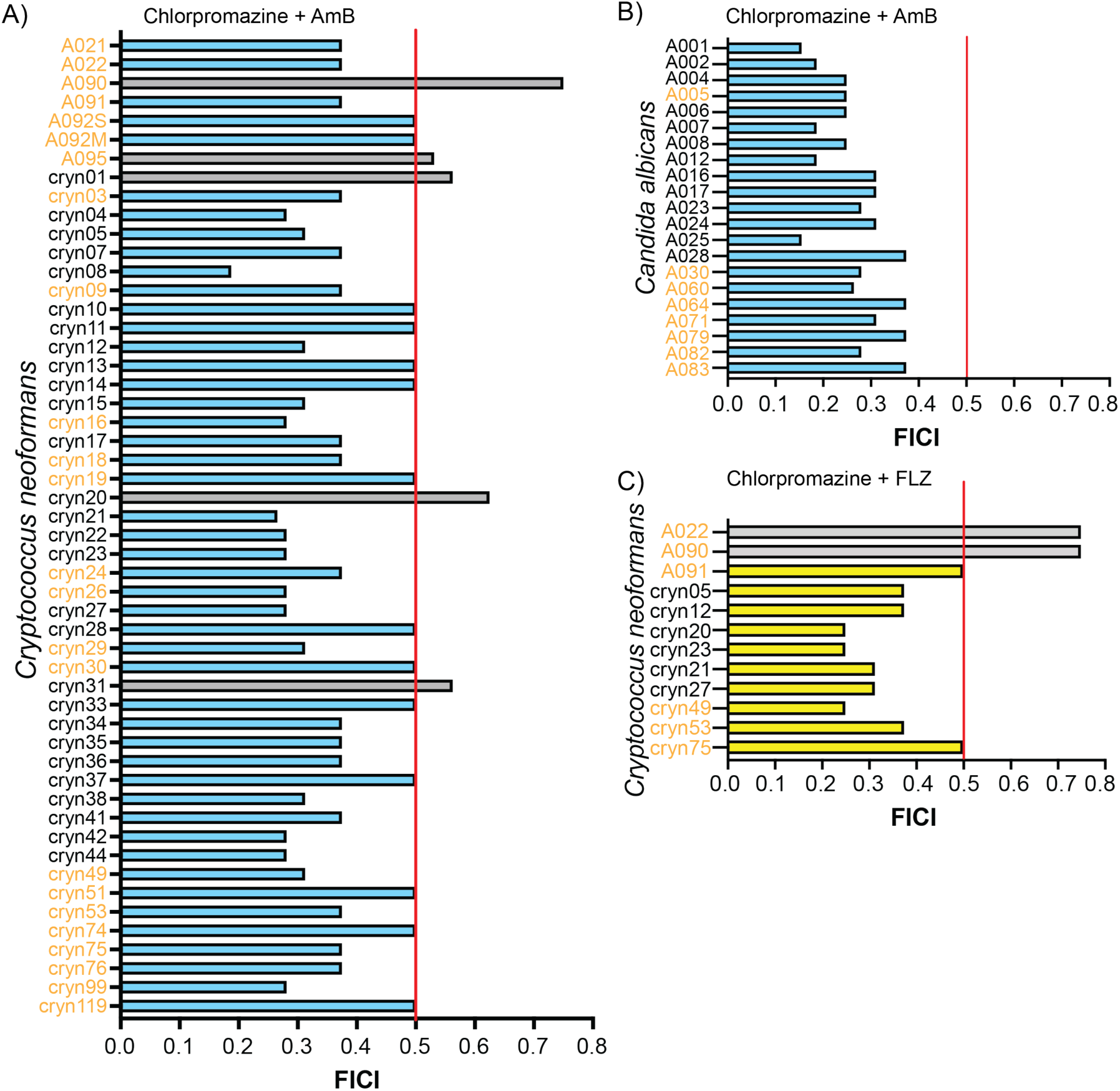
Synergistic effect of chlorpromazine in combination with amphotericin B or fluconazole in multiple clinical isolates and different fungal strains. A. *C. neoformans* synergistic interaction of chlorpromazine with AmB and B. fluconazole. C. Candida albicans clinical strains. Blue and green bars represent an FICI score of ≤ 0.5, indicating a synergistic result. Gray bars represent a FICI > 0.5, indicating no interaction. Red vertical line marks the division between synergistic and non-synergistic at the 0.5 mark. Names of strains resistant to fluconazole are labeled in orange.

**Table 1.** List of *C. neoformans* clinical isolates resistant and susceptible to fluconazole.

**Table 2.** FICI scores and MICs of *C. neoformans* treated with chlorpromazine and AmB combination. Non-synergistic are marked in blue.

Chlorpromazine in combination with AmB also acts synergistically against *Candida albicans* clinical isolates: all tested isolates, including 8 fluconazole-resistant isolates, showing a FICI of < 0.4 (**Fig. 1B** and **Table 3**). To assess whether chlorpromazine would exhibit similar synergistic activity when combined with fluconazole, we tested 12 *C. neoformans* isolates and 10 showed a synergistic response (**Fig. 1C** and **Table 4**). One isolate that did not show a synergistic response to chlorpromazine + fluconazole, A022, exhibited a synergistic response to chlorpromazine + AmB. Isolate cryn20 showed the reverse: a non-synergistic to chlorpromazine + AmB combination but a synergistic response to chlorpromazine + fluconazole (**Fig. 1C**).

**Table 3.** FICI scores and MICs of *Candida albicans* treated with chlorpromazine and AmB combination.

**Table 4.** FICI scores and MICs of *C. neoformans* treated with chlorpromazine and fluconazole combination. Non-synergistic are marked in blue.

Although the mechanism conferring fluconazole resistance in these clinical isolates is unknown, fluconazole resistance is commonly associated with mutations in Erg11 (46; 47; 48; 49), aneuploidy (50; 51; 52), and drug transporters (53; 54). AmB is an effective broad-spectrum antifungal against which resistance rarely (55) develops due to fitness tradeoffs (56). However, the emergence of *Candida auris* has developed compensatory mechanisms that make AmB-resistance more common (57), appearing in ∼18-30% of clinical isolates (58; 59). Polyene resistance involves modifications to the cell membrane and changes to the activation of cell stress responses (60). Only two of the fluconazole resistant *C. neoformans* clinical isolates tested did not show a synergistic response to AmB + chlorpromazine (A090 and A095). Taken together, these results show that chlorpromazine strongly inhibits fungal cell growth when combined with either AmB or fluconazole against multiple *C. neoformans* and Candida clinical isolates *in vitro*, including fluconazole-resistant strains.

### Chlorpromazine decreases lanosterol and affects transcript levels of genes involved in the ergosterol synthesis pathway

Ergosterol is an abundant sterol essential to fungi and its biosynthetic pathway is target of multiple antifungals (61). Sterols are synthesized in the endoplasmic reticulum (ER), then transported to the cell membranes to maintain membrane integrity (62). Due to chlorpromazine’s amphiphilic structure, it is hypothesized to insert into membrane bilayers by intercalating the hydrophobic head into the membranes (34). We first tested whether chlorpromazine treatment damages the plasma membrane. Neither chlorpromazine alone or AmB alone significantly increased permeability using propidium iodide, which cannot cross the plasma membrane of living cells (63). However, when both drugs were combined, the dye uptake, a measurement of increased permeability, increased 5-fold (**Fig. S1**).

We then considered that the increase in plasma membrane permeability could be an indirect result of chlorpromazine damaging other intracellular organelles, which affects the mechanism for membrane repair (64). Since sterols being necessary to regulate the plasma membrane permeability and fluidity (65), we focused on measuring the effects of chlorpromazine in ergosterol biosynthesis (**Fig. 2A**). We first examined the ergosterol biosynthesis pathway *ERG* genes’ transcript levels via RT-PCR. Late-stage ergosterol biosynthesis genes *ERG2*, *ERG4*, and *ERG6*, which are found in the late stage of the ergosterol pathway, significantly increased over control (**Fig. 2B**). Erg2, Erg4, and Erg6 proteins are predominantly located to the ER, where ergosterol is synthesized and distributed to intracellular and plasma membranes (66).

**Figure 2.**
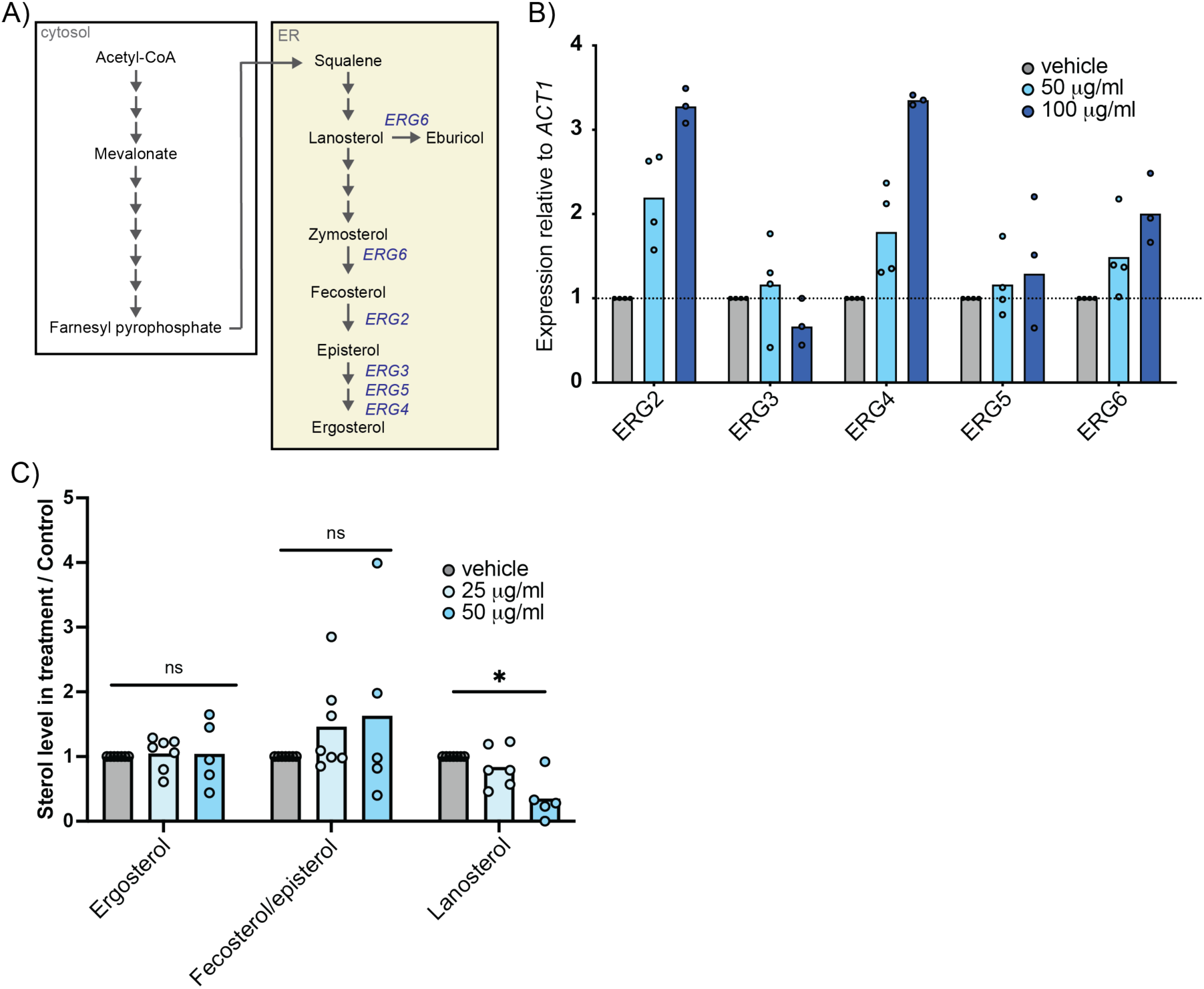
Chlorpromazine increases transcript levels of ERG2, ERG4, and ERG6 in chlorpromazine treated *C. neoformans*. A. Ergosterol biosynthesis pathway, including *ERG* genes in blue. B. Relative transcript levels normalized to control of cells treated with chlorpromazine. C. Sterol quantification of cells treated with increasing chlorpromazine concentrations. *Mann-Whitney p=0.026.

Next, we then investigated whether chlorpromazine directly affects ergosterol and its sterol intermediates along the pathway by measuring sterol levels via mass spectrometry. After a 24h treatment, we found that although ergosterol levels remained the same after cells were treated with chlorpromazine, lanosterol levels significantly decreased (**Fig. 2C**). Lanosterol is the first sterol in the sterol pathway after squalene is converted to squalene epoxide, an upstream precursor of the ergosterol biosynthesis pathway. This significant reduction in lanosterol levels suggests that chlorpromazine may interfere with the ergosterol biosynthesis pathway, which cells compensate for by upregulating genes late in the pathway to maintain ergosterol levels. Genes early in the pathway are not upregulated (Fig. SX).

The differential *ERG* gene regulation suggests that the response to chlorpromazine is complex and may involve sterol metabolism and possibly a stress response to membrane damage. Our data indicate that despite lower lanosterol levels, other sterol precursors may compensate for its reduction. It is possible that *ERG4* and *ERG6* which encode enzymes with low substrate specificity, can utilize alternative sterol substates for ergosterol synthesis. The increased expression of specific *ERG* genes without increasing *ERG3* and *ERG5*, suggests that chlorpromazine selectively influences steps in the pathway which can allow a better understanding on how *C. neoformans* responds to antifungal treatment and membrane damage.

### Fatty acid biosynthesis genes and lipid droplet formation are required to tolerate chlorpromazine

To identify chlorpromazine targets or targeted pathways, we screened two available (30; 67; 68; 69). *C. neoformans* single-gene knockout mutant library to search for sensitive or resistant mutants. We found 112 knockout mutants that showed sensitivity to chlorpromazine (**Table 5**). A metabolic enrichment pathway analysis using fungiDB (70) revealed significantly enriched pathways in fatty acid biosynthesis and degradation (p-value <0.05) and inositol lipids, all linked to processes that affect lipid composition (**Fig. 3**).

**Figure 3.**
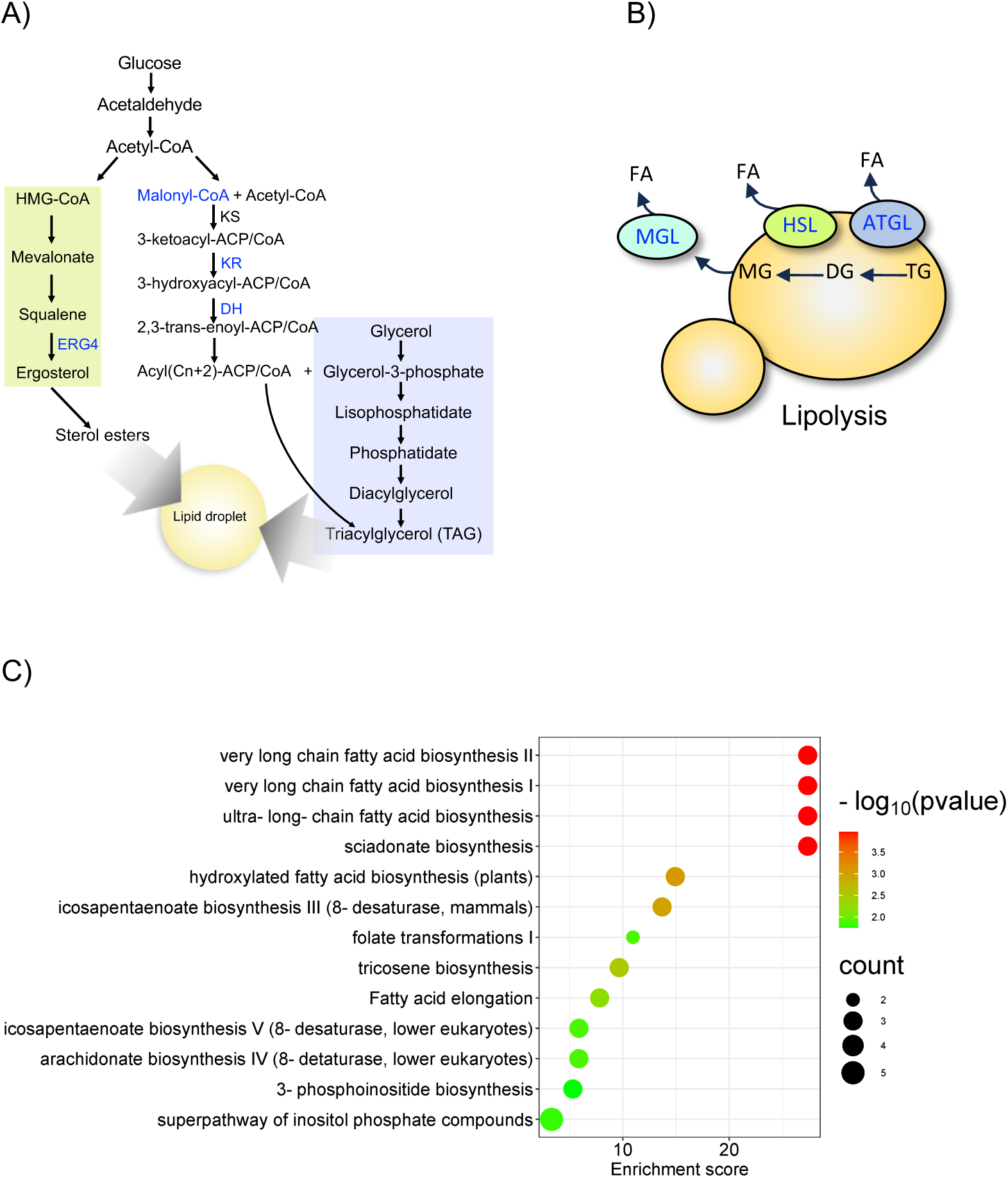
Metabolic enrichment pathway analysis revealed enriched pathways linked to processes that affect lipid composition. A) Fatty acid synthesis pathway and ergosterol biosynthesis pathway showing final products triacylglycerol (TAG) and sterol esters that are eventually stored in lipid droplets for slow release according to cell requirements. B) Schematic of degradation of lipid droplets via lipolysis. Knockout mutants that had increased sensitivity to chlorpromazine are marked in blue. C) SR plot created from a metabolic enrichment pathway analysis. FA: Fatty acid, MGL: monoacylglycerol lipase, HSL: hormone sensitive lipase, ATL: adipose triglyceride lipase. KS: 3-ketoacyl-ACP synthase. KR: 3-ketoacyl-ACP reductase. DH: 3-hydroxyacyl-ACP/CoA dehydrogenase. MAG: monoacylglyceride. DAG: diacylglyceride. TAG: triacylglyceride. G: glycerol.

**Table 5.** List of gene knockout mutants sensitive to chlorpromazine.

Lipids are categorized as fatty acyls (FA), glycerolipids (GL), glycerophospholipids (GP), sphingolipids (SP), sterol lipids (ST), prenol lipids (PR), saccarolipids (SL), and polyketides (PK) with each category further divided into specific classes and subclasses (71). Fatty acids, glycerolipids, glycerophospholipids, and sphingolipids all contain fatty acids as part of their structure. Fatty acids are not only sources of membrane-building components, but as previously mentioned, alterations in this metabolism leads to impaired cell function or death.

An important step in fatty acid regulation is the formation of lipid droplets. These are used for storage and gradual release of lipid species according to cellular demands for energy production, membrane synthesis, and lipid signaling (72; 73). Lipid droplets are organelles composed of an outer monolayer of phospholipids and proteins that mainly contain neutral lipids, such as triacylglycerols and sterol esters (72). Lipid droplets vary in size, from small and barely visible (0.2-0.5um) to large and prominent droplets (1um-2.5um) (74).

To investigate the involvement of chlorpromazine in the formation of lipid droplets, we used the fluorescent probe BODIPY to visualize and measure lipid droplet size and number in treated fungal cells. The area of lipid droplets significantly increased after treatment with chlorpromazine (**Fig. 4A**). Moreover, this is time-dependent: lipid droplet area significantly increased when cells were treated for 3h compared to 1h (p-value: 0.0037 Mann-Whitney), suggesting a dynamic lipid droplet change over time with drug treatment (**Fig. 4A**). Quantification of the number of lipid droplets per cell showed that chlorpromazine-treated cells also had fewer lipid droplets, with 30-50% of cells containing only 1 or 2 lipid droplets per cell (**Fig. 4B**). We occasionally noticed that chlorpromazine can surround lipid droplets (**Fig. S2**). DMSO (vehicle)-treated cells had a larger distribution in the number of lipid droplets, ranging from 1 to 20 lipid droplets per cell (**Fig. 4C**). These results suggest that chlorpromazine reduces the number of small lipid droplets to favor the formation of enlarged lipid droplets in *C. neoformans*. The disproportional enlargement of lipid droplets can have both beneficial and detrimental effects. Although the enlargement of lipid droplets can provide a storage capacity for excess lipids and protect from lipotoxicity, persistent lipid accumulation disrupts signaling pathways, membrane trafficking, and protein synthesis (75; 76; 77; 78).

**Figure 4.**
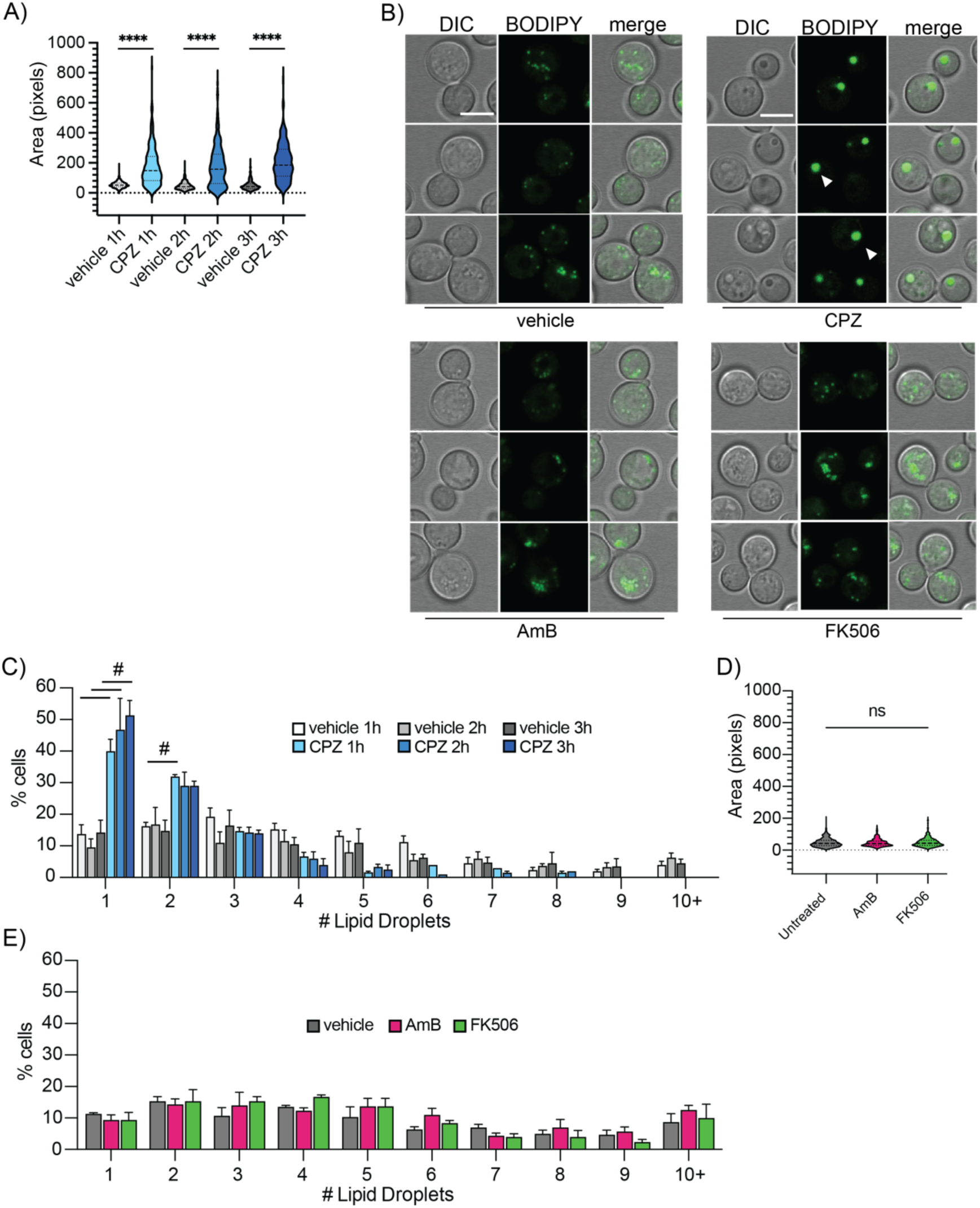
Chlorpromazine induces enlarged lipid droplets in *C. neoformans*. A) Measurement of area of lipid droplets after 1h, 2h, and 3h. of incubation with chlorpromazine. Area was quantified in relative pixel size. B) Confocal microscopy images of cells treated with DMSO control, 100µg/ml of chlorpromazine, amphotericin B, or FK506 stained with BODIPY. White arrows mark enlarged lipid droplets. Scale bar: 5 µm. C) Percentage of cells with a determined number of lipid droplets. X-axis shows the total number of lipid droplets per cell (1–20). D) Area measurement of lipid droplets from *C. neoformans* cells treated for 3h with AmB or calcineurin inhibitor FK506. E) Confocal microscopy images of cells treated with corresponding drug and stained with BODIPY. Scale bar: 5 µm C. Quantification of number of lipid droplets per cell. ****p<0.0001 We counted ≥100 cells per experiment and performed three independent experiments. Mann-Whitney ****p<0.0001. Unpaired t-test # p ≤ 0.05.

Lipid droplet formation is hypothesized to allow efficient storage of fatty acids during lipid overloading (73; 79). To verify that the formation of enlarged lipid droplets was specific to chlorpromazine, we treated fungal cells with 400 ng/ml FK506 or 100 ng/ml AmB (**Fig. 4B**). AmB and FK506, a known calcineurin inhibitor (80), are not known to have a direct effect on other fatty acids. Neither drug increased the area of the lipid droplets (**Fig. 4D**) and the number of lipid droplets per cell did not increase over control (**Fig. 4E**). This shows that stress unrelated to lipid synthesis does not induce changes to lipid droplet number of size and that the effect is specific to chlorpromazine. *C. neoformans* knockout mutants with roles in lipid biosynthesis that were sensitive to chlorpromazine showed morphological defects and an inability to form enlarged lipid droplets . Chlorpromazine also accumulated around lipid droplets (**Fig. S2**). It is possible that the formation of enlarged lipid droplets may help sequester chlorpromazine, potentially containing the drug and preventing further intracellular damage.

### Chlorpromazine targets vacuolar protein sorting genes

We additionally identified 8 knockout mutants that showed resistance to chlorpromazine that belong to the vacuolar protein sorting process and are involved in ESCRT pathway functions: *vps20*τι, *vps22*τι*, vps23*τι, *vps25*τι*, vps27*τι*, vps28*τι*, vps32*τι*, vps36*τι (**Fig. 5B**). Others have reported that lipid droplet utilization is coupled to organelle homeostasis and ESCRT function (81; 82). *VPS* mutants negatively affect lipid droplet consumption, resulting in reduced efficiency in utilizing or consuming lipid droplets (83). However, lipid droplets still increased in size in *VPS* mutants when treated with chlorpromazine (data not shown).

**Figure 5.**
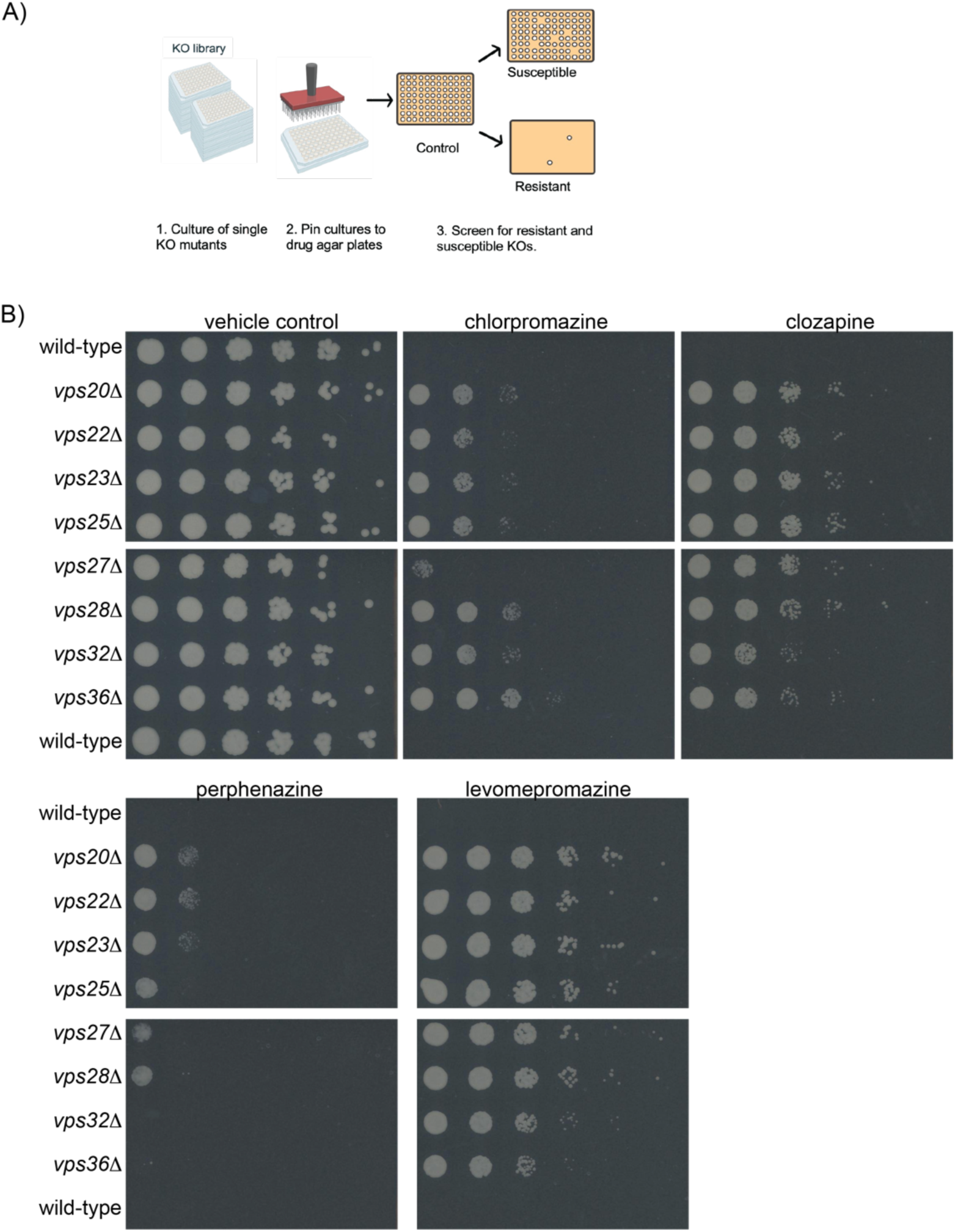
*Vps* knockout mutants are resistant to chlorpromazine and analogs. A) Schematic of screening workflow. B) Wild-type (KN99) and mutant cells grown in the presence of chlorpromazine, clozapine, perphenazine, or levomepromazine. Consecutive spots represent a serial 5-fold dilution of wild-type or knockout mutant cells. Images from three days of growth at 30°C but were taken one days 2-4. Mutant cells’ resistance to these molecules is visible on days 2-4.

### Chlorpromazine analogs act synergistically with fluconazole and AmB

While we have previously tested synergistic combinations in mouse infection models (23), chlorpromazine treatment in rodents present several challenges that made finding an appropriate dose that did not cause harm difficult (data not shown). Specifically, chlorpromazine affects cellular immune responses (84; 85), causing agranulocytosis (86), loss of alveolar macrophages in rats (87), and suppression of IL-6 in monocytes. Given these confounding factors, testing chlorpromazine itself in mice in the context of infection is likely to result in data that are difficult to interpret.

Instead, we hypothesized that drugs with similar chemical structure to chlorpromazine can have similar effects against *C. neoformans*. We chose three different chlorpromazine analogs with varying chemical substructure similarity: clozapine, perphenazine, and levomepromazine. Clozapine is less likely to cause motor side effects (88) and perphenazine (89) and levomepromazine (90) exhibit lower incidence of immunomodulatory side effects. Perphenazine and levomepromazine are first-generation antipsychotics. Clozapine is a second-generation antipsychotic known for its higher effectiveness and reduced adverse side effects compared to other atypical antipsychotics and chlorpromazine (91; 92).

We used ChemMine tools (93) to calculate similarity using the Maximum Common Substructure (MCS) Tanimoto score (**Fig. 6A**). The Tanimoto coefficient is a standard metric that measures distance or molecular similarity and ranges between 0 to 1 with values closer to 1 indicating higher similarity (94). The analogs clozapine, perphenazine, and levomepromazine had a MCS Tanimoto score of 0.42, 0.77, and 0.83, respectively.

**Figure 6:**
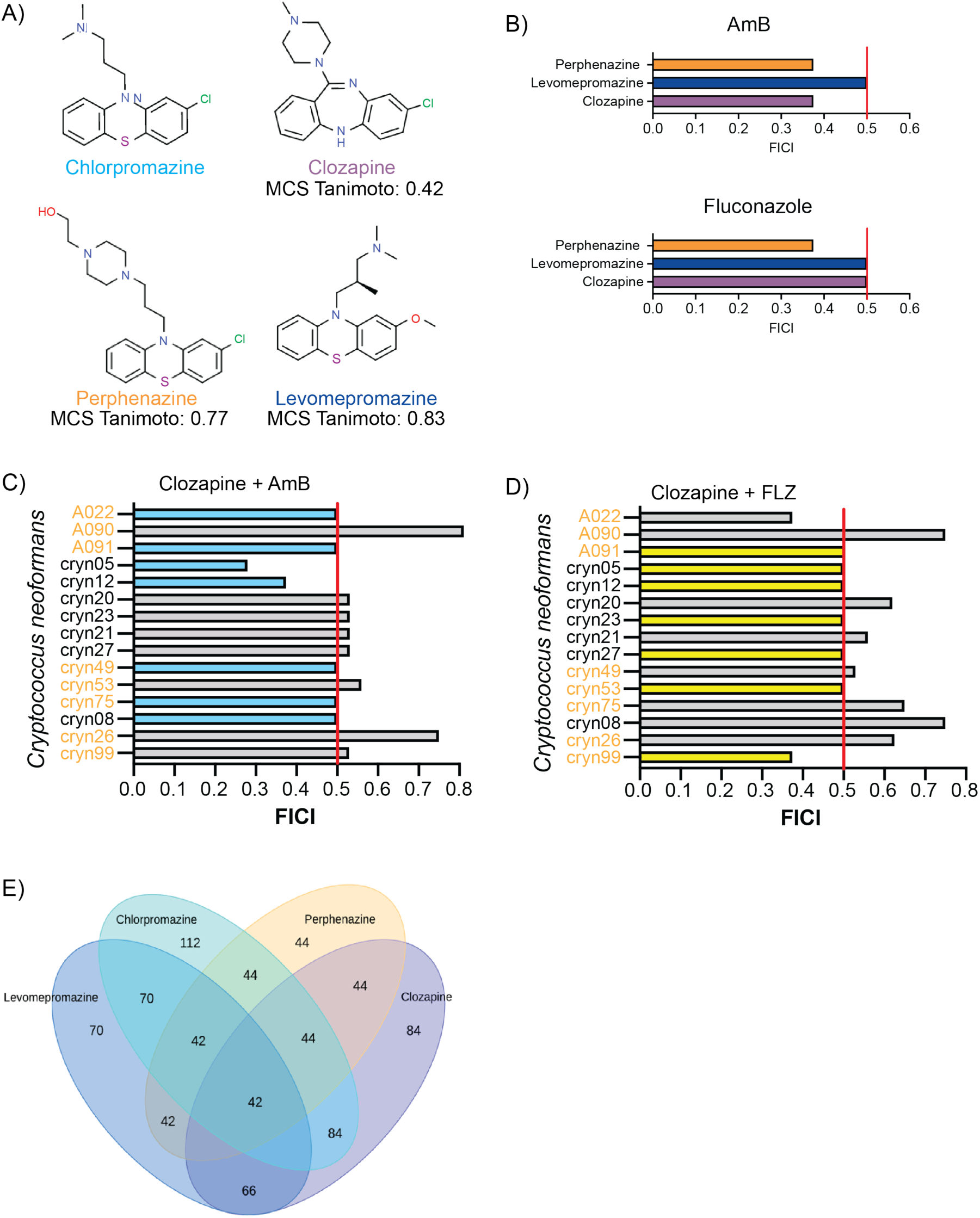
Chlorpromazine analogs act synergistically with AmB and fluconazole. A) Chlorpromazine analogs clozapine, perphenazine, and levomepromazine with different degree of structural similarity B) Synergistic FICI scores of the different analogs in combination with AmB (Top) or fluconazole (bottom). FICI scores of clozapine in combination with C) AmB and D) Fluconazole. Blue (AmB) or yellow (fluconazole) bars represent an FICI score of ≤ 0.5, indicating a synergistic result. Gray bars represent a FICI > 0.5, indicating no interaction. Red vertical line marks the division between synergistic and non-synergistic at the 0.5 mark. Names of strains resistant to fluconazole are labeled in orange. E) Venn diagram showing the overlap of single-gene knockout mutants sensitive to chlorpromazine and its three analogs. The numbers below each analog represent the subset of mutants that were sensitive to the respective drug among the mutants sensitive to chlorpromazine. Chlorpromazine sensitive mutants were identified from an entire single-gene knockout mutant library screen.

When we tested the three analogs for synergy with AmB and fluconazole against our lab strain KN99, all three chlorpromazine analogs acted synergistically with both antifungals regardless of varying structural similarity (**Fig. 6B**) with a FICI of :: 0.5. We further tested clozapine in combination with AmB or fluconazole against *C. neoformans* clinical isolates and both combinations, respectively, showed synergy in 46% and 53% of the isolates tested (**Fig. 6C,D**).

Single gene knockout mutants resistant to chlorpromazine are similarly resistant to chlorpromazine analogs. We further investigated whether the same single-gene knockout mutants that were sensitive to chlorpromazine had similar growth when exposed to clozapine, perphenazine, and levomepromazine. Only 44 of the 112 knockout mutants sensitive to chlorpromazine were also sensitive to perphenazine. In comparison, clozapine and levomepromazine induced sensitivity in 84% and 70% of the knockout mutants respectively (**Fig. 6E**).

## Discussion

Synergistic drug combinations are promising potential treatments for systemic fungal infections. They have the capability to overcome resistance (**Fig. 1**), lower treatment costs, and delay the development of drug resistance (29; 95). This study aims to elucidate the molecular mechanisms underlying the synergistic interactions between AmB and the antipsychotic chlorpromazine, revealing a potent combination with enhanced antifungal efficacy. Moreover, chlorpromazine’s less toxic analogs, perphenazine, levomepromazine, and clozapine appear to act via the same mechanism and are themselves promising antifungal treatments.

Previous research showed that chlorpromazine could interact with cellular membranes in different organisms and cell types (96). The effects of this disruption extend from membrane dissolving in very high concentrations to increased membrane permeability with lower drug concentrations. Others showed that chlorpromazine has a reduced affinity to membranes that contain higher cholesterol levels, the analog of ergosterol in mammalian cells (97). Here, we saw that AmB increases lipid droplet size, a response to stress that phenocopies nutrient starvation (72) and allows regulated breakdown of lipid droplets to prevent lipotoxicity (98) to mitochondria and other organelle. In combination with AmB, chlorpromazine effectively disrupts membranes, increasing membrane permeability 5-fold (**Fig. S1**). However, with lipids sequestered in large lipid droplets, the cell might not be able to repair the plasma membrane, resulting in severe cell damage (**Fig. 7**). It is possible that in *C. neoformans,* the reduction in ergosterol availability when treated with AmB increases the affinity of chlorpromazine to membranes. This idea is also supported by the fact that chlorpromazine synergizes with fluconazole, a drug known to reduce ergosterol in fungi (**Fig. 1B**) (99).

**Figure 7.**
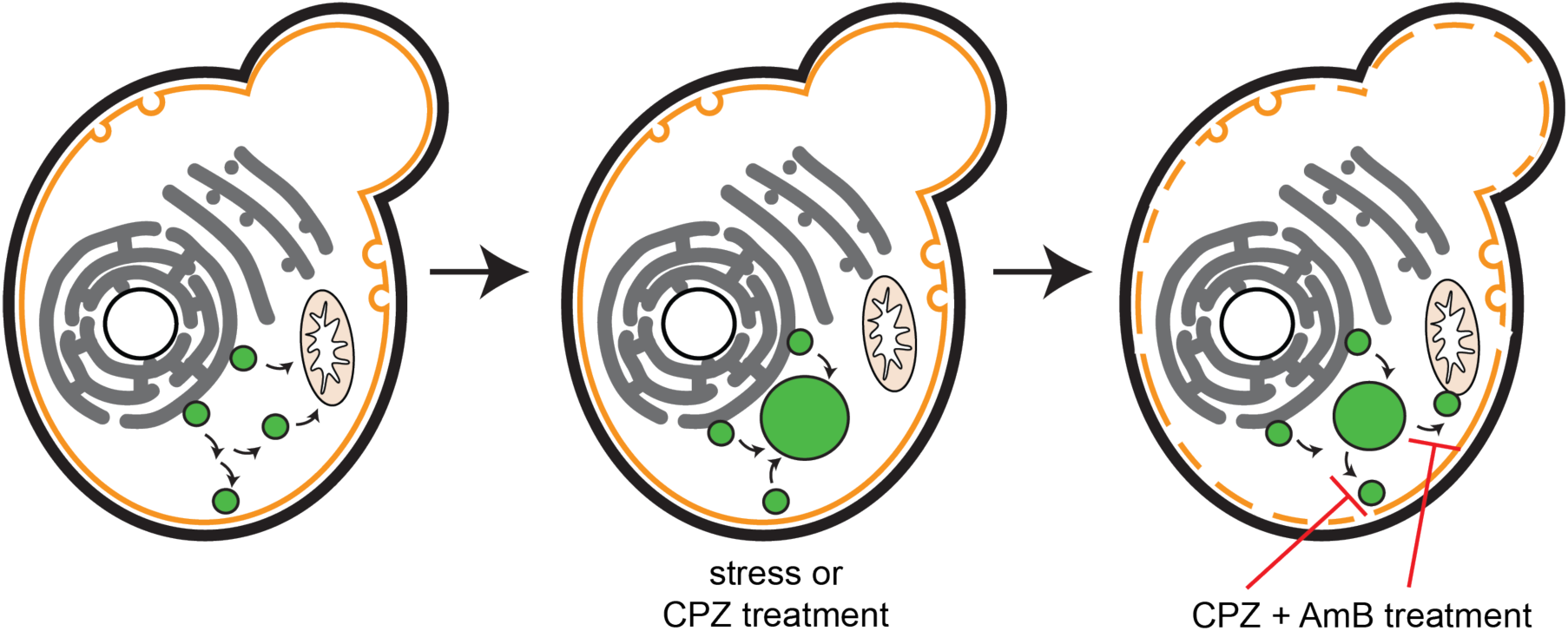
Chlorpromazine + AmB increase membrane permeability and chlorpromazine increases lipid sequestration in lipid droplets. Larger lipid droplets can slow response to sterol stress, rendering cells treated with chlorpromazine more sensitive to AmB.

Chlorpromazine is not the only psychiatric drug with antifungal properties: the selective serotonin reuptake inhibitor sertraline also inhibits fungal cell growth (100; 101). Sertraline can also increase lipid droplet size in *C. neoformans* (102). The authors proposed that the increase in the size of lipid droplets was due to a surplus of phosphatidic acid known in *Saccharomyces cerevisiae* to induce fusion events in lipid droplets. When there is an increase in the number of lipid droplets in cells, droplet fusion events increase, favoring the eventual coalescence of lipid droplets (73). We hypothesize that this occurs when chlorpromazine-treated cells show enlarged lipid droplets (Fig. 4).

We established that chlorpromazine has a broad efficacy against multiple fungi *in vitro*. However, further investigation is needed to reduce and address potential side effects when testing *in vivo*. The related side effects specific to chlorpromazine include benign leukopenia and, on rare occasions agranulocytosis. Different studies have indicated that chlorpromazine exhibits diverse immunomodulatory effects. The IL-6 suppression is particularly relevant to AIDS patients, a group highly susceptible to cryptococcal infection, since IL-6 was shown to directly upregulate HIV replication (103). In a study testing chlorpromazine toxicity in rats, the drug showed reduced 5-week survival and loss of alveolar macrophages (87). Future studies are needed to clarify how this drug affects the immune cell profile in mice during a cryptococcal infection.

Chlorpromazine analogs are promising potential antifungal drugs. We demonstrated that chlorpromazine analogs in clinical use show synergy with AmB and fluconazole. Clozapine, a second-generation antipsychotic, has a lower risk of certain adverse effects compared to chlorpromazine. Future research can further optimize these analogs for enhanced efficacy and safety profiles against fungal infections. These results highlight the potential of repurposing antipsychotic analogs for the development of novel treatments for fungal infections.

## Acknowledgments

This work was supported by grant R01AI137331 from NIH/NIAID to J.C.S.B. J.M.B was supported by T32AI055434 from NIH/NIAID and C.T.M. was supported by T32GM141848 from NIH/NIGMS.

## Methods

### *C. neoformans* growth and small-molecule MIC determination

MIC assays were performed with 1X YNB + 2% glucose. Overnight cultures were grown at 37°C and diluted to OD600 = 0.0292. Small molecules were dissolved in DMSO at varying stock concentrations, and serially diluted in a 2-fold series in a 96-well plate. Each well was inoculated and plates were incubated for 48 hours at 30°C. MIC values were calculated by measuring OD600 using the Molecular Devices SpectraMax iD5 plate reader. OD600 difference between 48h and 0h of above 0.1 is considered the line for MIC90.

### Checkerboard assays

Cells were grown overnight and adjusted to OD_600_ = 0.0292 as starting inoculum for all species of fungi. Drugs were serially diluted in 96-well plates as described previously (23), and all wells were inoculated with 2 µL of cells. Plates were incubated in 30°C with shaking/resuspension at 24 and 48 hours. OD_600_ for each plate was measured at 0 and 48 hours with a Molecular Devices SpectraMax iD5 plate reader for growth inhibition assessment. FICI scores were calculated as described in Wambaugh *et al.* (104). Briefly, FICI scores are determined by a 90% growth inhibition cutoff for MICs using this equation:

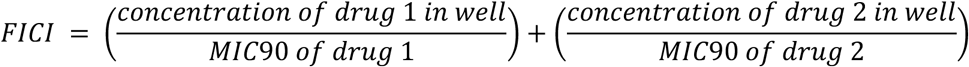

FICI scores of ≤0.5 are considered synergistic. Checkerboard assays are repeated at least twice for each strain and drug combination. If the result (synergistic or not synergistic) is the same, the average FICI is shown on graphs. If the result is not the same (e.g. one synergistic, one not), the assay is repeated a minimum of five times and the scores from the predominant result (i.e. all scores over 0.5 or all scores ≤ 0.5) are averaged and shown on the graph.

### RT-PCR

Quantification of ergosterol biosynthesis pathway gene expression intermediates was done by RT-PCR. *C. neoformans* cells were treated overnight with each drug. 1×10^7^ cells were pelleted, flash frozen in liquid nitrogen, and lyophilized until the pellet was completely dry. RNA was extracted using the Qiagen RNeasy column kit (Catalog no. 74014) with modifications.

Pellet was resuspended in 600µl of RTL buffer and approximately 50µl of 1mm and 150µl of 0.5mm zirconium beads were added to each sample. Samples were placed in the BioSpec mini bead beater to lyse cells with 12x 2 min pulsations, placing blocks and samples on ice for 5 min in between pulses to prevent overheating. RNA was then extracted following kit instructions. cDNA was synthesized using random primers (Invitrogen), M-MLV reverse transcriptase (Invitrogen), 10x dNTPs and incubating at 37°C for 1h. cDNA was further purified using Zymo DNA clean and concentrator (Cat. no. 11303) following kit instructions. Real-Time qPCR was performed using the Applied Biosystems QuantStudio 3 system and SYBR green for continuous fluorescent monitoring. C_t_ values were obtained in triplicate, averaged and normalized to ACT1 calculating relative gene expression using the 2^−ΔΔCt^ method.

### Knockout mutant library drug agar screening

Drug agar plates were made by adding small molecules dissolved in DMSO into 2X YNB + 4% glucose and mixing a 50:50 ratio of drug/YNB and agar for a final concentration of 2% agar. Single KO mutant collections were constructed by the Hiten Madhani group in the University of California, San Francisco and the Yong-Sun Bahn group at Yonsei University - both purchased from the Fungal Genetics Stock Center. Frozen stocks of the mutants were pinned onto YPAD agar to grow at 30°C and liquid cultures were made in 1X YNB + 2% agar. Liquid cultures were pinned onto drug agar plates using a 96-well replicator pinner. Plates were then incubated at 30°C and images were taken every day for 4-5 days using a flatbed scanner. Resistant and susceptible mutants were identified by eye.

### Small molecule sensitivity testing

*C. neoformans* strain WT or single-gene knockout mutants obtained from the Madhani collection (30) were streaked onto YPAD agar, grown at 30°C for at least 48 hours. Liquid cultures were made in 1X YNB + 2% glucose and incubated overnight. Fungal cells were adjusted to an initial concentration of 1×10^7^ cells/mL and 5-fold serially diluted across six wells in a 96-well plate. 3 µL of each dilution was pipetted using a multichannel pipette onto each plate containing drug agar (300ug/ml CPZ and 450ug/ml clozapine) or DMSO control. Agar plates were incubated at 30°C, and images were taken daily for 4-5 days using a flatbed scanner. The spot assay was repeated twice and representative images were shown.

### Confocal microscopy with BODIPY staining

Liquid cultures of *C. neoformans* strain KN99 and knockout mutants were grown in YPD media at 37°C overnight. Fungal cells were adjusted to 3×10^6^ cells/mL and treated with the appropriate drug for 3 hours (100 µg/mL clozapine, 100 µg/mL CPZ, 100 ng/mL AmB, or 400 ng/mL FK506) then centrifuged and washed with 1X YPD. For chlorpromazine time course experiments, cells were treated with 100 µg/mL of CPZ for 1, 2, and 3 hours. Cells were then treated with BODIPY stain in 500 µL YPD for 30 minutes. BODIPY stain was dissolved from powder with anhydrous DMSO into a 10 mM solution and adjusted to 1 mM. BODIPY solution was added to each sample to achieve a final concentration of 3uM. After staining, the cells were washed twice with 1X PBS and resuspended in 1X PBS for microscopy.

Cells were prepared for microscopy in a MatTek glass bottom dish. Images were taken using the Zeiss LSM 880 Airyscan confocal microscope with a 63X objective lens. Within the Zeiss Zen image acquisition software, the BODIPY FL (488 nm laser) and DAPI (405 nm laser) channel settings were used, as well as T-PMT for DIC.

### Lipid droplet analysis

The CellProfiler software (105) was used to analyze images of BODIPY stained fungal cells. A pipeline was created to quantify the number of cells, area of each lipid droplet, and number of lipid droplets per fungal cell in a total of 100 cells per sample. The area was defined by the number pixels within a lipid droplet. The number of lipid droplets per cell was determined by defining lipid droplets that were contained within each parent cell as “child objects”. Three independent biological replicates were used.

---

**Supplemental Figure 1.**
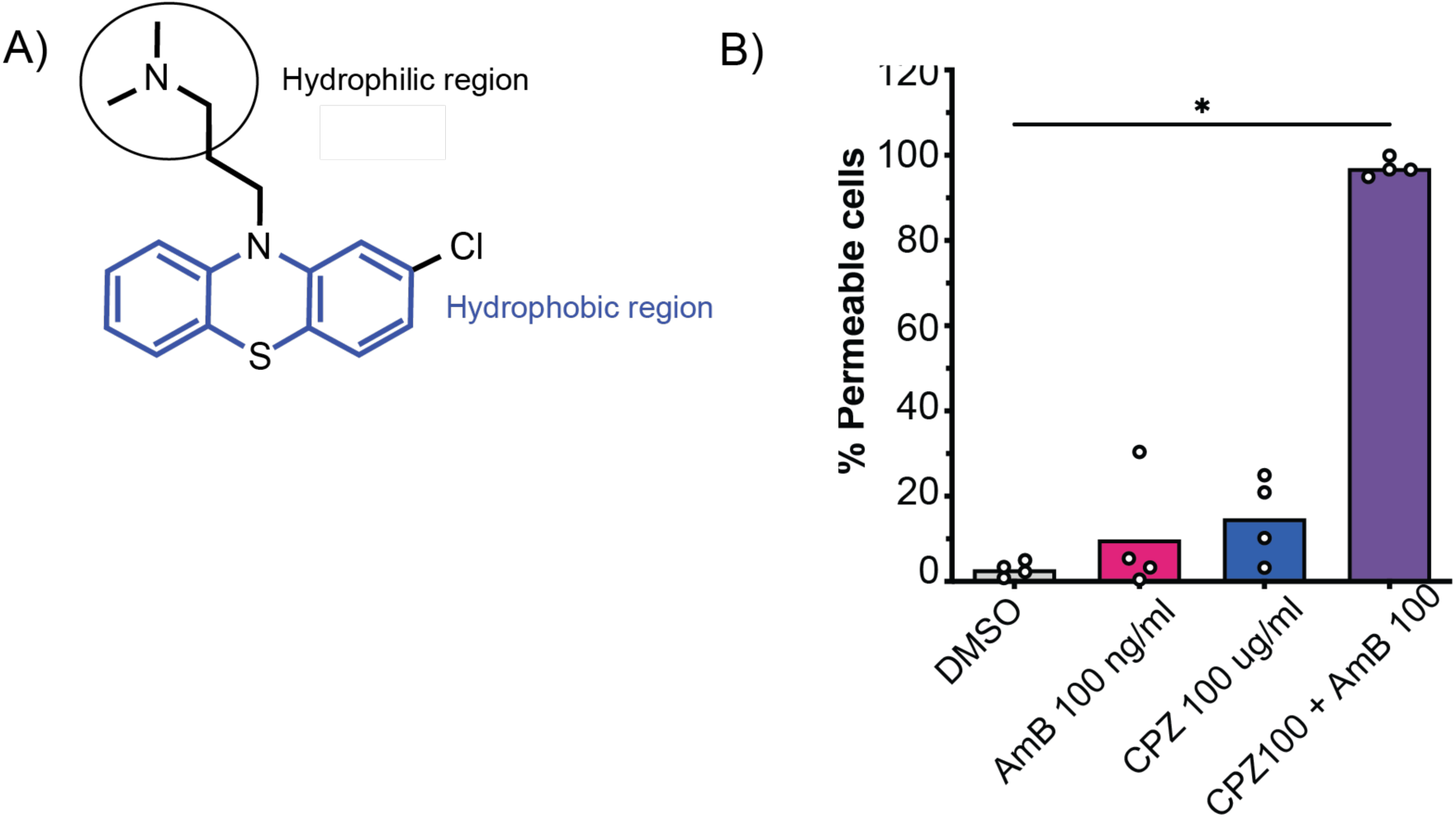
Chlorpromazine in combination with AmB increases membrane permeability in *C. neoformans*. A) Chlorpromazine molecular structure showing hydrophilic (Circle) and hydrophobic (blue) regions. B) Percent of membrane permeability measured by flow cytometry in cells treated for 3h with AmB, chlorpromazine, or combination and stained with propidium iodide (PI). *Mann-Whitney p=0.0286

**Figure S2:**
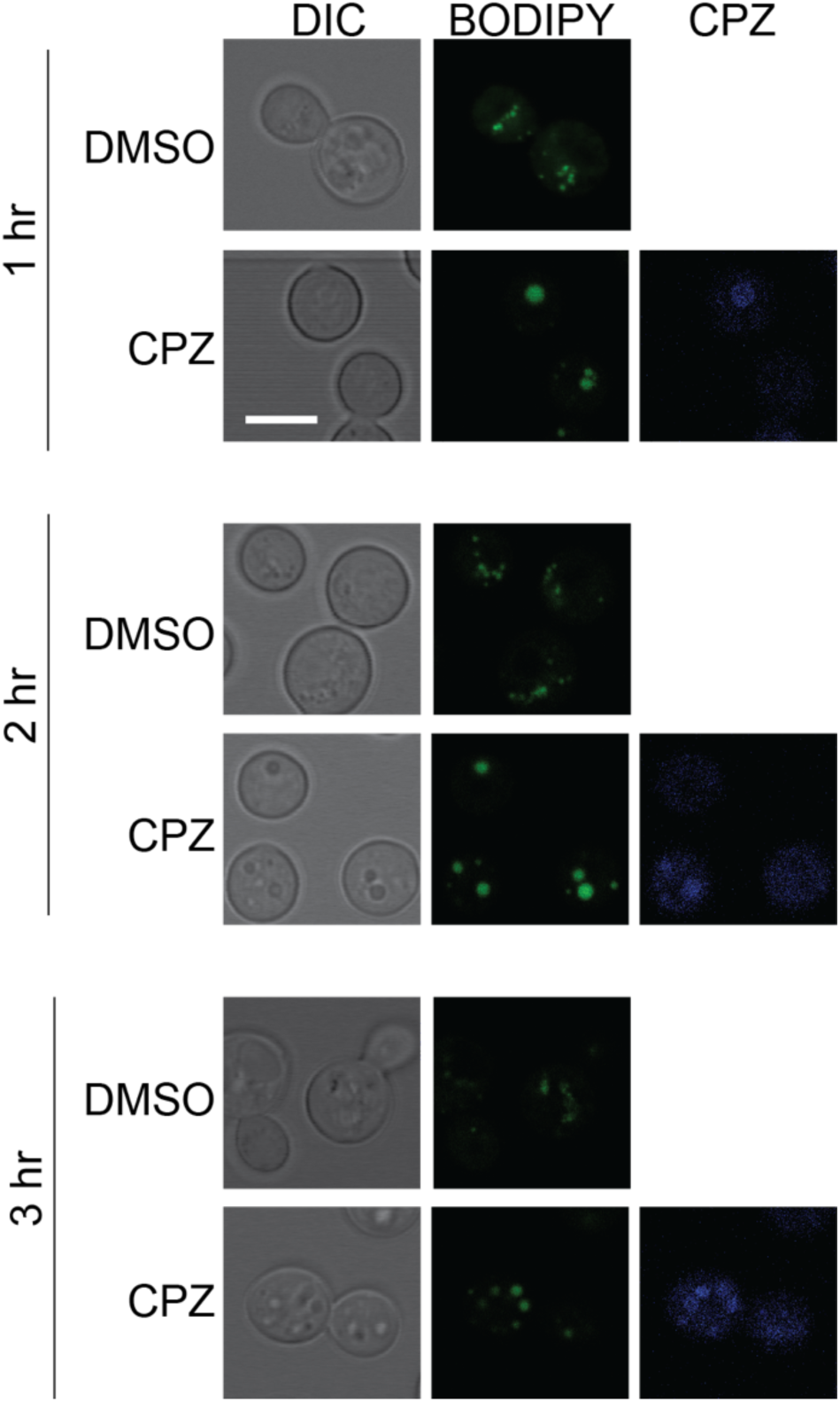
Chlorpromazine accumulates around lipid droplets. Cells treated with either vehicle control (DMSO) or chlorpromazine (CPZ) for 1, 2, or 3 hours show accumulation of naturally fluorescent CPZ around the lipid droplets (BODIPY). Scale bar represents 5 μ

